# Functional and epitope specific monoclonal antibody discovery directly from immune sera using cryoEM

**DOI:** 10.1101/2024.12.06.627063

**Authors:** James A. Ferguson, Sai Sundar Rajan Raghavan, Garazi Peña Alzua, Disha Bhavsar, Jiachen Huang, Alesandra J. Rodriguez, Jonathan L. Torres, Maria Bottermann, Julianna Han, Florian Krammer, Facundo D. Batista, Andrew B. Ward

## Abstract

Antibodies are crucial therapeutics, comprising a significant portion of approved drugs due to their safety and clinical efficacy. Traditional antibody discovery methods are labor-intensive, limiting scalability and high-throughput analysis. Here, we improved upon our streamlined approach combining structural analysis and bioinformatics to infer heavy and light chain sequences from electron potential maps of serum-derived polyclonal antibodies (pAbs) bound to antigens. Using ModelAngelo, an automated structure-building tool, we accelerated pAb sequence determination and identified sequence matches in B cell repertoires via ModelAngelo derived Hidden Markov Models (HMMs) associated with pAb structures. Benchmarking against results from a non-human primate HIV vaccine trial, our pipeline reduced analysis time from weeks to under a day with higher precision. Validation with murine immune sera from influenza vaccination revealed multiple protective antibodies. This workflow enhances antibody discovery, enabling faster, more accurate mapping of polyclonal responses with broad applications in vaccine development and therapeutic antibody discovery.

## Introduction

As of 2024, antibodies comprised 206 of globally approved therapeutics, reflecting their growing efficacy in treating diverse diseases(*1*). In 2018, antibodies represented 20% of US Food and Drug Administration (FDA) approved therapeutics(*2*). Their safety in humans(*3*) and rapid clinical translation make them an attractive approach to conditions like cancer, autoimmunity, and infectious disease(*4*, *5*). Initially discovered using hybridoma technology(*6*), the invention of phage display in the 1990s(*7*), followed by improvements using yeast(*8*) or mammalian displays(*9*), have significantly enhanced antibody discovery throughput. Single cell B-cell receptor (BCR) sequencing with or without antigen specific sorting enables a high throughput means of isolating monoclonal antibodies (mAbs) to a specific target as well as providing insights into the diversity and evolution of immune responses(*10*). However, identifying epitope specific mAbs remains resource-intensive, requiring extensive screening through binding and functional assays (*11*, *12*). These bottlenecks highlight the need for alternative approaches to identify epitope-specific mAbs more efficiently particularly through advancements in BCR-sequencing data analysis(*17*, *18*). Such innovations could streamline the mAb discovery process and accelerate the identification of protective epitopes and antibody responses.

Our laboratory’s development of cryoEMPEM (cryoElectron Microscopy Polyclonal Epitope Mapping) for characterizing polyclonal antibody (pAb) responses was a significant step forward in being able to assess which epitopes are targeted by an individual immune response to vaccination or infection(*19–21*). High-resolution information contained in the pAb regions of cryoEMPEM maps can also be used to determine Ab sequences from the structure (SFS), enabling the matching of epitope specific antibodies to BCR-sequencing data sets(*22*). While the initial study successfully determined two epitope-specific mAbs by combining cryoEM and next generation sequencing (NGS) data, limitations remained, particularly the time intensive map interpretation. Additionally, the isolated mAbs showed poor expression yields and targeted non-functional epitopes on the envelope glycoprotein of human immunodeficiency virus 1 (HIV-1). Recognizing the need for efficiency and the removal of human bias, we have now integrated the AI automated model builder, ModelAngelo (MA)(*23*) and it’s built in HMMER search capabilities(*24*), into our SFS workflow which significantly enhances the speed and accuracy of both model building and structural analysis.

Here, we outline a method for integrating MA in conjunction with pAb electron potential maps and BCR-sequencing to derive epitope specific antibody sequences. First, we benchmarked MA’s capabilities with the two polyclonal maps that were previously used to develop the manual SFS method as part of a non-human primate(NHP) HIV-1 Env vaccination trial(*22*). Antibodies synthesized based on the MA and subsequent HMMER search recommendations exhibit higher yields with comparable binding affinity compared to those developed through conventional methods. We then conducted a mouse vaccination study with the neuraminidase (NA) glycoprotein of influenza virus combined with paired BCR-sequencing to test the capability of MA to determine Ag-specific pAb sequences with a new dataset. Ultimately, we produced five NA inhibiting mAbs that were functional in vivo and protected against viral challenge. The increased speed and accuracy of this new approach greatly expands the efficiency and power of antibody discovery using cryoEMPEM.

## Results

### Benchmarking ModelAngelo for cryoEMPEM SFS

To benchmark MA as a pAb building tool we ran its build_no_seq mode with the maps from the first SFS dataset from NHPs vaccinated with soluble BG505 HIV-1 Envelope glycoprotein with the SOSIP stabilizing mutations(*25*). This dataset included two cryoEMPEM maps of pAbs – Rh.33104 pAbC-1 (pAbC-1) and Rh.33172 pAbC-2(pAbC-2). pAbC-1 targets the glycan hole and pAbC-2 targets a strain specific epitope in the V2/V3 region, as was confirmed by the original SFS paper(*22*). For both cryoEM maps, MA generated potential pAb models and .hmm files (Figure 1A, S1). While MA built complete fragment variable (fv) regions of the polyclonal Fab (pFab) pAbC-1, pAbC-2 models were fragmented in both heavy and light chains(Figure S1). For both pAb models, the amino acid assignments did not accurately represent an antibody sequence, with an average sequence identity of 58.9% to the Abs sequences confirmed in the original SFS paper (Figure S2). However, MA generates a hidden Markov model (.hmm) file for each chain constructed – visualized as a logoplot in Figure 1A generated by Skylign(*26*). To generate a .hmm for the heavy and light fv of the fragmented pAbC-2 we generated auxiliary code that would merge fragmented .hmm files together, using a pAb aligned polyalanine fv model to assess which .hmm files to merge for the heavy and light chains.

**Figure 1.**
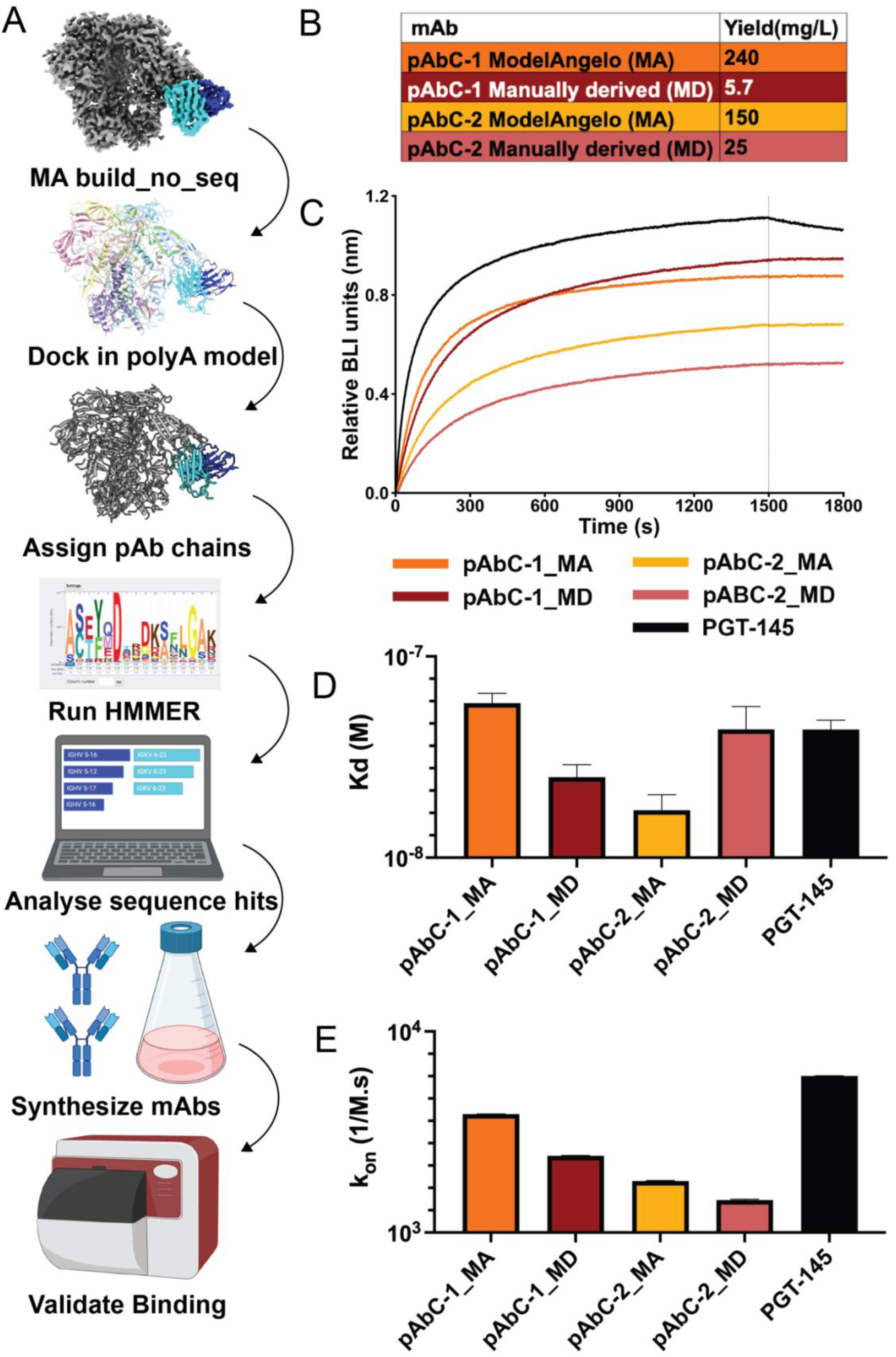
(A) MA-workflow. Starting with an electron potential map (Density corresponding to the antigen shown in grey, heavy chain in blue, and light chain in cyan) ModelAngelo is run in build_no_seq mode. Chains and the subsequent HMMs are assigned as heavy and light chains. A representative HMM is shown as a logoplot generated by Skylign. Heavy and light chain sequence matches are searched for in BCR-sequencing repertoire using .hmm files generated by MA using the MA inbuilt HMMER. The top scoring hits are synthesized for binding validation. (B) Comparison of antibody yields for manually and MA derived antibody sequences. (C) Representative BLI curves for highest concentration of HIV-1 Env (1000nM). (D) Steady state affinities for manually and MA derived pAbC-1 and pAbC-2 antibodies. (E) On-rates for manually and MA derived pAbC-1 and pAbC-2 antibodies. MA, model angelo; MD, manually derived.

Using the heavy and light chain .hmm files that we generated from our maps, we ran MA’s HMMER search on the corresponding B-cell repertoire for epitope specific heavy and light chains. This identified new putative heavy and light chain sequences with an average 92.3% identity to the original manually derived SFS mAbs (Figure S3). We then expressed these new Abs to validate binding.

Before considering affinity, we observed that MA derived antibodies showed higher yields than the manually derived sequences (Figure 1B). For the human manually derived mAbs, pAbC-1_HU and pAbC-2_HU, we got yields of 5.7 mg/L and 25 mg/L respectively. Whereas for the MA-derived mAbs, pAbC-1_MA and pAbC-2_MA, the yields were 240 mg/L and 150 mg/L respectively. We measured affinities by biolayer interferometry (BLI) and since no measurable off rate could be determined (Figure 1C, S4), for all mAbs we compared the steady state affinities and on rates (Figure 1D and E, respectively). For pAbC-1 the manually derived mAb had a marginally higher affinity with an apparent steady state Kd of 25 nM compared to the MA-derived mAb with 59 nM. However, for pAbC-2 the human derived mAb had a Kd of 43nM which was a lower affinity than the MA-derived pAbC-2 with Kd of 17 nM. When looking at the on-rates the situation is reversed. For pAbC-1, the MA-derived mAb had an on rate of 3.9 x10^-3^ M^-1^s^-1^ which was higher than the human-derived mAb with an on rate of 2.4 x10^-3^ M^-1^s^-1^. For pAbC-2 the manually derived mAb bound faster with on rate of 1.8 x10^-3^ M^-1^s^-1^ compared to the MA-derived mAb at 1.4 x10^-3^ M^-1^s^-1^. These results highlight that MA can rapidly identify epitope-specific antibodies with higher yields and comparable binding profiles in under 2 hours.

### Testing ModelAngelo on an influenza virus neuraminidase vaccination study in mice

With the method successfully benchmarked, we tested the workflow to discover functional influenza antibodies from mice vaccinated with NA from A/Indiana/10/2011 (Ind11) H3N2 (Figure 2A). CryoEMPEM analysis on sera pooled from 10 mice revealed robust antibody responses against multiple NA epitopes (Figure S5). Because we were interested in functional antibodies, we specifically generated a 3.3Å polyclonal electron potential map with a pAb binding the active site of NA with our processing workflow shown in Figure S5 and data collection statistics in Figure S6. We ran MA build_no_seq on this map to build an initial model and .hmm files and, as with the pAbC-2 benchmarking map, the MA model of the active site pAb was fragmented. Likely due to the polyclonality and multi-animal pooled sera of the active site binding pAb responses, the cryoEM reconstruction of the pAb CDR regions was ambiguous so we inserted 3 additional positions with random amino acid probabilities into the CDRH3 as well as 2 in the CDRH1 and 2 in the CDRH2 regions in the heavy chain .hmm.

**Figure 2.**
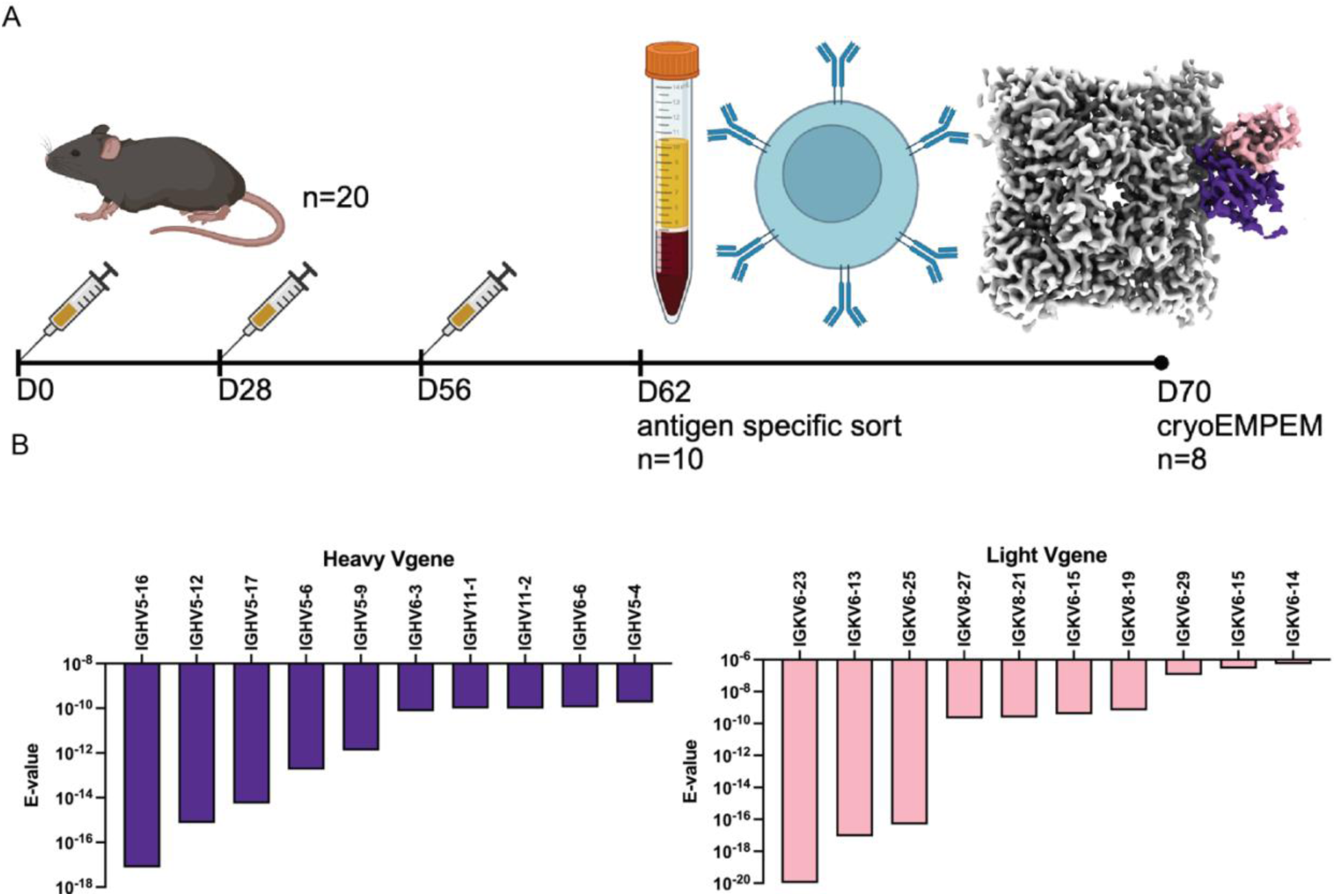
(A) Vaccination scheme for cryoEMPEM and B cell sorting. Density corresponding to the heavy and light chains of the active site pAb are shown in purple and pink respectively. The NA density is shown in white. (B) Top gene usages for the heavy and light chains of the active site region pAb.

For this batch of 10 mice, we were unable to obtain BCR sequences. In their absence, we used the V gene germline database for C57BL/6 mice from the International Immunogenetics system (IMGT)(*27*) as an input for HMMER search to determine the gene usage of the pAb (Figure 2B). The HMMER E-value was used to rank the gene usage. Based on the E-values, the top five most likely IGHV gene usages were IGHV5-16, IGHV5-12, IGHV5-17, IGHV5-6, and IGHV5-9. For the light chain, the top 5 most likely IGLV or IGKV gene usages were IGKV6-23, IGKV6-13, IGKV6-25, IGLK8-27, and IGLK8-21.

Another 8 mice underwent the same vaccination scheme and PBMCs were harvested to generate a database of roughly 5000 NA-specific paired antibody sequences for our HMMER search. We selected 4 mAbs using the heavy chain .hmm and 3 mAbs using the light chain .hmm. For the heavy chain .hmm, mAbs were selected based on the lowest E-scores and the top four V gene usages: IGHV5-16, IGHV5-12, IGHV5-17, and IGHV5-6 (Figure 2B, 3A). For the light chain we chose three sequences with the lowest E-scores, all using the most likely light chain gene IGKV6-23, but with three different heavy chain genes (Figure 3A). Six of the 7 selected mAbs bound to Ind11 NA by ELISA (Figure 3B), and 5 inhibited recombinant Ind11 NA using an enzyme-linked immunosorbent assay (ELLA) (Figure 3C). Before conducting protection studies, we confirmed that these mAbs inhibited NA in the context of live Ind11 influenza virus (Figure 3D).

**Figure 3.**
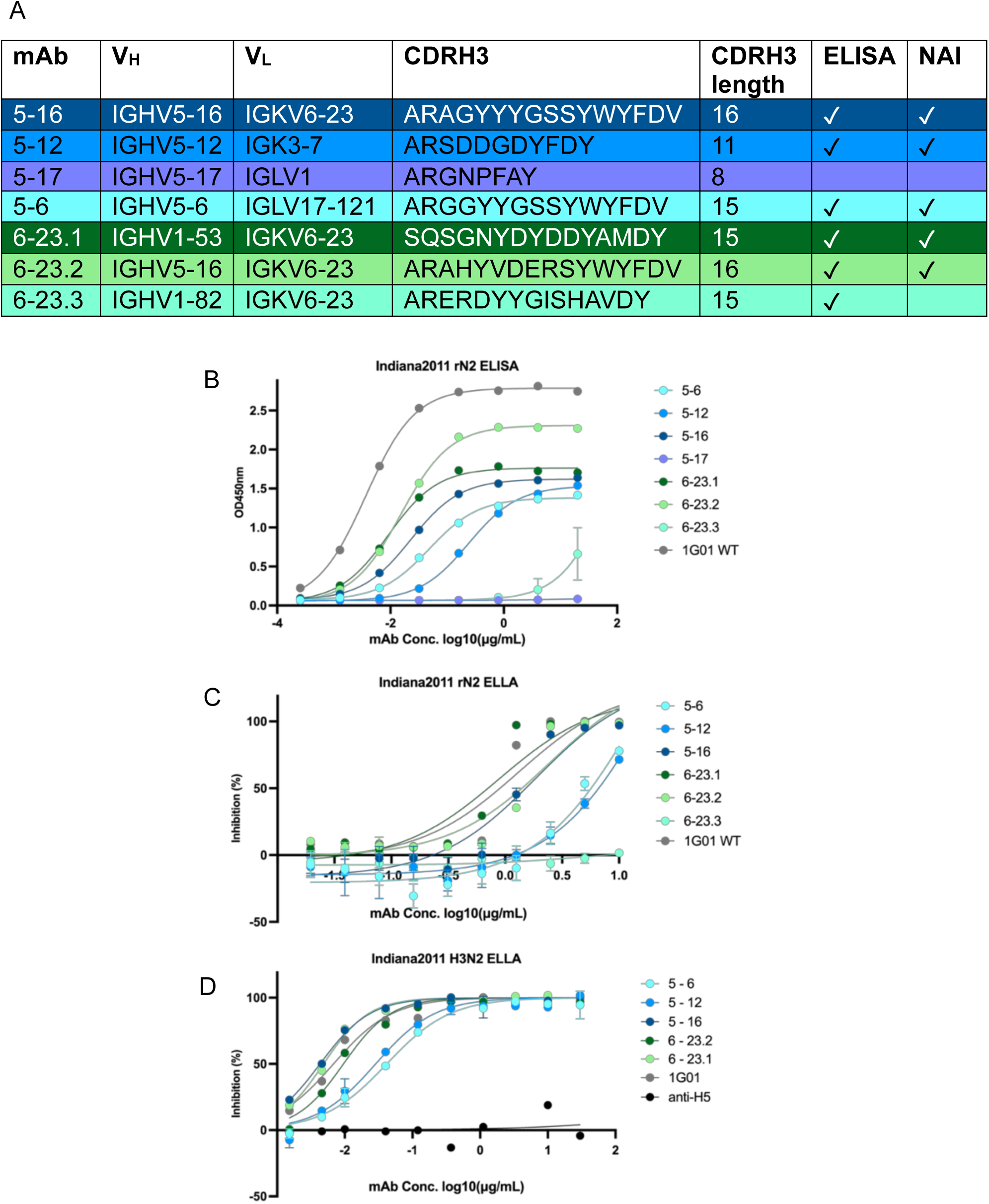
(A) Table of selected mAbs from HMMER search. (B) ELISA of selected mAbs against Ind11 NA. (C) ELLA of selected mAbs against recombinant Ind11 NA. (D) ELLA of selected mAbs against live Ind11 influenza virus.

### Structural epitope analysis of NA-inhibiting mAbs

To map the epitopes targeted by the five NA inhibiting mAbs, we solved cryo-EM structures of these mAbs in complex with the Ind11 NA, with resolutions ranging from 2.8Å to 3.4Å (Figure 4A, S6), and validation metrics and CDRH3 densities shown in Figure S7. To compare the isolated mAbs with the pAb we built a representative pAb model using the germline IGHV5-16 and IGHJ1 sequences with a CDRH3 length of 16, and light chain using the germline IGKV6-23 and IGKJ3. We used Ablang2(*28*) to predict the missing CDRH3 residues that would have been inserted during VDJ recombination, then built the pAb model using Abodybuilder2(*29*), followed by manual fitting in COOT(*30*) with the NA model.

**Figure 4.**
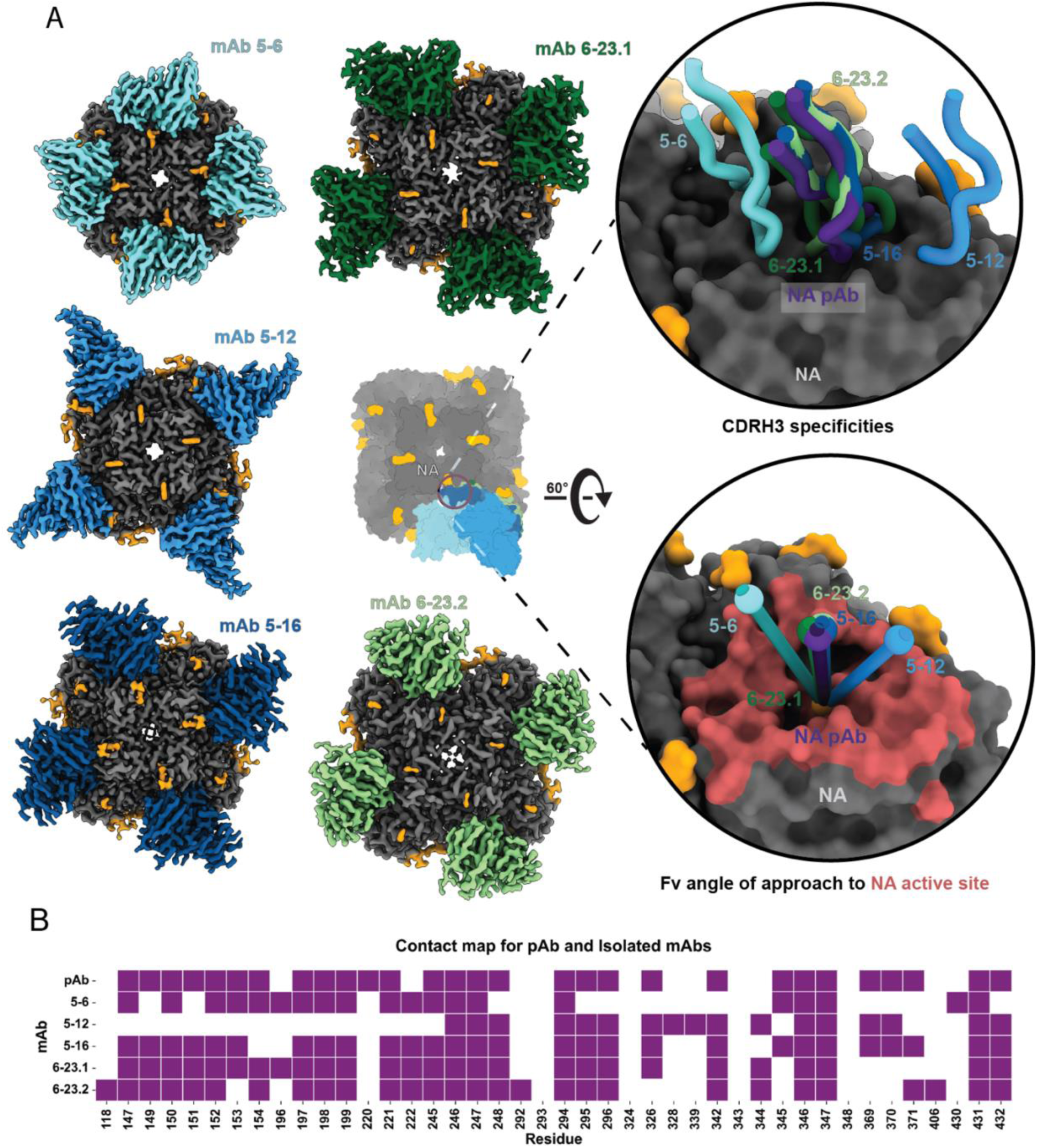
(A) Electron potential maps of the 5 isolated mAbs including a peanut model overlay of the mAbs. The top inset of the active site compares the backbones of the CDRH3 regions. The bottom inset compares the angle of approach of each mAb. The angle is measured between the mAb centroid, the active site centroid, and the pAb centroid. (B) Residues that contributed more than 5Å^2^ of BSA due to mAb or pAb binding. The list of residues listed were determined using the getcontacts interface analyzer tool.

Using the protein interface analyzer tool, getcontacts(*31*), we determined all NA contact residues utilized collectively by the pAb and the five mAbs, shown in red on the NA surface in Figure 4A and listed in the contact plot in Figure 4B. To assess similarity, we compared the buried surface area (BSA) of each of the NA contact residues for each mAb to the pAb using Biopython(*32*), normalizing against unbound NA. Residues contributing more than 5Å BSA are shown in Figure 4B with a normalized BSA heatmap in Figure S8A. These results indicated that that 5-16 is most structurally similar to the pAb, differing by 3 contact residues, while 5-12 differed by 17 of 42 collective residues. Pearson correlation coefficients for normalized BSA (Figure S8B) suggest 5-16 (0.93) and 6-23.1 (0.89) most closely resemble the pAb. Root mean square deviation (RMSD) Cα after model alignment to the pAb bound protomer, showed 5-16,5-6, and 6-23.2 has the lowest Cα RMSDs (1.7Å), while 6-23.1 and 5-12 had an RMSD of 2.7Å. Lastly, we compared the angle of approach of each mAb compared to the pAb (Figure 4A). Angle of approach calculations revealed that all mAbs had less than 3° difference from the pAb except, for 5-12 and 5-6 which had differed by 28°. Overall, our structures suggest that 5-16 and 6-23.2 are the most structurally representative of the active site pAb that we identified in terms of contacts, BSA contribution, and RMSD Cα, and these mAbs also exhibit the great inhibition potency to live virus.

### Protection study in mice

To evaluate whether the generated mAbs using the developed method could inhibit viral replication *in vivo*, we tested the different antibodies in a prophylactic setting. 6-week-old DBA/2J mice were administered with 5 mg/mL of the mAbs 2 hours prior to intranasal infection with Ind11 H3N2 virus at a dose of 5 times the lethal dose 50% (LD_50_). Mice were then monitored for weight loss and survival for 14 days to assess protection (Figure 5). The antibodies with greatest neuraminidase inhibitory activity *in vitro* (5-16, 6-23.1, 6-23.2) also showed better protection from infection in mice, with those treated with 6-23.2 achieving complete survival similar to mice administered with positive control mAb 1G01, while mice receiving 5-16 and 6-23.1 had survival rates of 80% and 66.7% respectively. This degree of protection was not seen in mice administered with 5-12 or with the irrelevant anti-H5 mAb, indicating that the developed tool can predict and facilitate the generation of mAbs with effective protective capacity.

**Figure 5.**
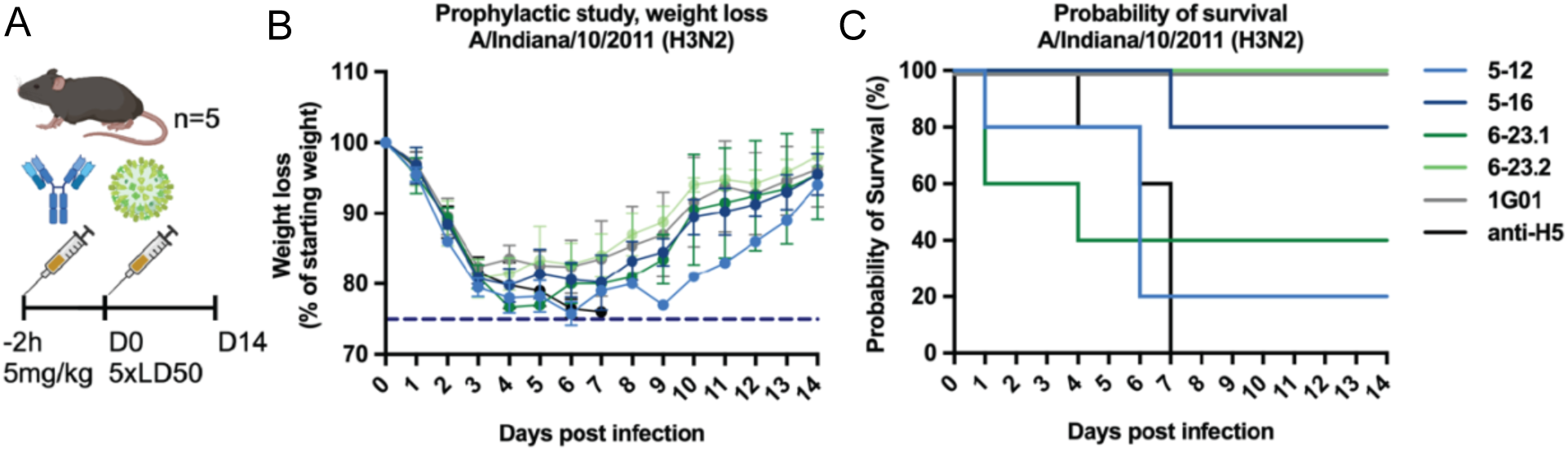
(A) Six-week-old female DBA/2J mice (n=5 mice per group) were injected intraperitoneally with 5 mg/kg of generated mAbs (5-12, 5-16, 6-23.1, 6-23.2), a well-defined anti-NA antibody was used as a positive control (1G01), and an irrelevant antibody (anti-H5), was used as negative control. The antibodies were administered 2 hours before the mice were infected with 5 times LD_50_ (lethal dose 50%) of the A/Indiana/10/2011 (H3N2) virus. The percentage of initial body weight (B) and survival (C) were monitored and plotted over a 14-day period post-infection for each antibody group. The percentage of body weight is relative to the starting body weight, and the data is represented as group means with standard deviation. (C)1G01 data is offset by 1% for clarity.

## Discussion

As was demonstrated in our original SFS manuscript and subsequent studies (*19*, *21*, *22*), the single particle nature of cryoEM enables the effective examination of the heterogeneous immune response to vaccination and infection. Extracting sequence information from pAb electron potential is a powerful immunological tool for identifying and epitope-specific antibodies within sequence repertoires(*22*). The integration of MA into this workflow has enhanced this pipeline as demonstrated well with our benchmarking dataset, and with more recent analyses(*33*, *34*).

Firstly, it has sped up the analysis of cryoEMPEM maps to under 2 hours meaning that data analysis is no longer a bottleneck. Secondly, unlike our previous manual SFS algorithm, MA generates .hmm files for faster, unbiased, probabilistic sequence searches, although having to manually merge .hmm files together and determine potential missing residues introduces some human bias. Despite this introduced bias we, along with others(*35*), observe that working with single small .hmm files from fragments of an fv, the best result achievable from a single incomplete .hmm is typically a V gene assignment. Lastly, using HMMER for exhaustive, bias-free searches has transformed the SFS pipeline into a high-throughput scalable tool for epitope specific antibody discovery.

The reduction in human bias likely contributed to the improved yields for the HIV-1 benchmarking pAbs, notably pAbC-1. The unpaired BCR-sequencing repertoires for the NHPs makes selecting a pair of functional sequences challenging. For pAbC-1, the MA-derived heavy chain had only two amino acid substitutions occurring in the framework regions. However, the light chain had 18 differences half of which occurred in framework region two. Given the location of these substitutions, it suggests that a large part of the yield improvements stem from improved mAb stability particularly in the heavy light chain interface.

Regardless of whether SFS is done manually or with MA a major limitation remains the quality of the electron potential map. Ambiguities arise from three main sources: sample specific issues leading to poor resolution of orientation bias, inherent flexibility of the pAb constant domains causing locally lower resolutions, and lastly, the polyclonal nature of pAb maps meaning that maps represent a composition of amino acids. Each of these pathologies can result in MA generating fragmented chains that we saw in pAbC-2 as well as our NA immunization dataset. The need to merge .hmm files and identify missing residues between chains, introduces human bias and extra analysis time. In the future it may be possible to use other probabilistic tools to determine these ambiguous regions. For example, inverse folding tools such as Antifold(*36*) or ProteinMPNN(*37*), with retrained weights(*38*), or fine-tined antibody language models(*28*) may be used in combination with initial protein structure models to predict probabilistic residues in regions deemed to be ambiguous. Even fine tuning of the MA model itself on only Ag:Ab structures may improve the accuracy of resultant .hmm files.

In our NA immunization dataset, we introduced 3 random amino acids in the CDRH3, and 2 in both CDRH1 and CDRH2, in our heavy .hmm to address polyclonality. Despite adding ambiguity, we discovered five functional mAbs all targeting the same epitope, with two that were able to inhibit as potently as the best in class mAb 1G01 (Figure 3D)(*39*). As postulated in the original SFS paper the complete sequence of the pAb may not needed to find functional antibodies. Notably, two of the five mAbs were discovered using only the light chain sequence alone – thanks to the paired BCR sequencing dataset. 6.23-1, discovered via the light chain, used the unexpected IGHV1-53, contrary to V gene usages shown in Figure 2. The most potent inhibitor, 5-16, closely resembled the pAb in the polyclonal map, a trend that continued for the remainder of the isolated mAbs with the exception of 6-23.1 and 6-23.2 where the former is more potent but slightly less similar to the pAb. The trend further follows with the protection data with 6-23.2 protecting 100%, 5-16 protecting 80%, and 6-23.1 protecting 66%. 5-12 which structurally diverges from the pAb the most only protected 25% of the mice and 5-6 did not protect at all, however, for this mAb we could not attain a statistically significant result. While SFS is not a quantitative method, these findings demonstrate how cryoEMPEM captures the most abundant and tightest binding pAbs. However, despite the large difference in epitope footprint between 5-12 and 5-6 to the pAb the fact that the HMMER search suggested them as epitope specific mAbs implies that similar molecular features are being used by different clonotypes to target the same or overlapping epitopes on NA.

CryoEM is a valuable tool for understanding the structure function relationship of protective mAbs and is now a pivotal tool in the understanding of serum immune responses through characterizing pAb sequences from electron potential maps. Its single particle nature could allow for synergistic integration with quantitative techniques such as proteomics-based method Ig-seq(*40*, *41*). It may also be useful in discovering public clonotypes as in demonstrated in this study, where not only were the mice that were sequenced not the same mice that the pAb map came from, the pAb map also represents the composite response of 10 mice. Further, the cryoEMPEM maps enable us to focus in on serum mAbs to specific functional epitopes, such as the NA active site, which contrasts with hybridoma or antigen specific B-cell sorting strategies that generate antibodies in an epitope agnostic manner. Hence, the efficiency of on-target antibody discovery is thereby increased.

Machine learning tools will without doubt continue to improve the speed and quality of immune repertoire analysis which is a vital part of structure-based antibody and vaccine design(*42*). Here we show how the ML tool ModelAngelo has revolutionized our SFS workflow, rapidly deconvoluting pAb responses with speed and precision. This integration of ML into cryoEM-based immune profiling not only accelerates vaccine development but also enhances our ability to respond swiftly to emerging pathogens. By streamlining the discovery of antigen-reactive and protective antibodies, this pipeline paves the way for faster, epitope specific antibody identification in the face of future global health threats.

## Materials and Methods

### Expression and purification of BG505 SOSIP and PGT145

Antigen expression and purification were performed as described previously (17). Briefly, BG505 SOSIP.664 was expressed in HEK293F cells (ThermoFisher Scientific). The proteins were purified from cell supernatants using PGT145 immunoaffinity chromatography (PMID: 25589637). MgCl_2_ buffer (3 M) was used for protein elution from the immunoaffinity matrix. BG505 SOSIP.664 samples were concentrated, buffer exchanged to tris-buffered saline (TBS) (Alfa Aesar), and subjected to size exclusion chromatography (SEC). HiLoad 16/600 Superdex 200 pg (GE Healthcare) running TBS buffer was used for SEC purification. Fractions corresponding to the BG505 SOSIP.664 trimer were pooled, concentrated to 1 mg/mL, and frozen for storage.

PGT145 IgG was expressed in HEK293F cells (ThermoFisher Scientific) by co-transfecting plasmids containing IgG heavy chain, light chain, and TPST2 (tyrosylprotein sulfotransferase). Recombinant IgG was purified from cell supernatants using a prepacked MabSelect protein A affinity column (Cytiva), eluted with 0.2 M glycine pH 2.0, buffer exchanged to TBS buffer, and frozen.

### Expression and purification of N2 A/Indiana/10/2011 neuraminidase antigen

A/Indiana/10/2011 N2 was expressed by BioMetas using a modified version of a protocol previously described here(*43*). Briefly, NA was expressed in Expi293 cells using a Twin-Strep-Thrombin-hVASP-NA-ectodomain construct. Proteins were purified from cell supernatants using TwinStep column, concentrated and purified further using a SEC column. Fractions corresponding to NA were pooled, concentrated, and frozen for storage.

### Immunization experiments

8–10-week-old C57BL/6J mice (Jackson Laboratories, RRID:IMSR_JAX:000664) were immunized intraperitoneally with 10µg N2 (A/Indiana/10/2011, Biortus Biosciences Co. Ltd, BioMetas) formulated in Sigma Adjuvant System (Sigma-Aldrich) and boosted on days 28 and 56. 8 mice were terminated on day 70 for spleen collection and 10 mice on day 62 for submandibular blood collection into BD MicroTainer Blood Collection Tubes (Beckton-Dickinson). Serum was collected after centrifugation (5 min, room temperature, 20,000 x g). All immunization experiments were performed under Harvard University and Massachusetts General Hospital (MGH) International Animal Care and Use Committee (IACUC) approved protocols (2016N000286) in AAALAC-accredited MGH facilities.

### BCR Sequencing:Fluorescence-activated cell sorting

Mono-biotinylated N2 (A/Indiana/10/2011, BioMetas) was incubated with streptavidin (Streptavidin-AF647 and Streptavidin-PE, BioLegend) in a 4:1 ratio at least 30 min before staining. Splenocytes were enriched for activated B cells. Enriched splenocytes were stained in 50 nM probe mix (assembled as described above) for 30 min, followed by staining with fluorescent antibody mix (CD38-BV510, CD138-BV605, IgD-BV786, B220-FITC, CD95-PE-Cy7, NK1.1-APCeF780, Gr-1-APCeF780, F4/80-APCeF780, CD4-APCeF780, CD8-APCeF780), TotalSeq-C antibody mix (TotalSeq™-C0914 - CD273, TotalSeq™-C0557 - CD38, TotalSeq™-C0917 – CD95, TotalSeq™-C0810 – CD138, TotalSeq™-C0571 – IgD) and TotalSeq C hashtags (BioLegend) for 30 min. Cells were resuspended in phosphate buffered saline(PBS) + 2% fetal bovine serum(FBS) 200 ng/mL 4′,6-diamidino-2-phenylindole (DAPI) (Thermo Fisher Scientific). Cell sorting was performed on a BD FACSAria Fusion (BD Biosciences).

### BCR Sequencing: Cell encapsulation and sequencing

Cell encapsulation was performed using a Chromium Controller (10x Genomics) and the Chromium Next GEM Single Cell 5’ Kit v2. Libraries were built according to the manufacturer’s instructions. Libraries were sequenced using a NextSeq2000 Sequencing System (Illumina) as previously described(*44*).

### BCR Sequencing: Single-cell RNAseq analysis

Sample pre-processing was performed using the Cell Ranger analysis pipeline (v7.2.0, 10x Genomics) aligning to the mm10-2020-A reference transcriptome. Sample demultiplexing, QC, normalization and clustering were performed with the R package Seurat v4.3.0(*45*).

### Isolation of mouse sera IgG and Fab preparation for cryoEMPEM

EMPEM protocols were performed as described by the EMPEM with deviations described below (*46*). Sera was heat inactivated in a 56°C water bath for 60 min and filtered with a 0.2 um filter and Triton X-100 was added to a final concentration of 1% v/v. Total IgG was purified using a Cytiva ALIAS autosampler and Cytiva AKTA pure system using a pre-packed HiTrap MabSelect PrismA column (Cytiva). IgG was eluted with by incubation with 0.1M glycine pH 3.0 directly into wells containing 1M Tris-HCl pH 8.0 buffer. Samples were buffer exchanged into PBS using centrifugation with 30kDa cut-off Amicon concentrators. Total IgG was digested with papain in freshly prepared digestion buffer (20mM sodium phosphate, 10 mM Ethylenediaminetetraacetic acid(EDTA), 20 mM cysteine, pH 7.4) at 37°C for 5 hours and was quenched with excess iodoacetamide. Samples were buffer exchanged into PBS using 10kDa cut-off Amicon concentrators and Fc fragments were removed with Capture select resin. To purify any further serum impurities and papain, the polyclonal fab mixture was run through size exclusion chromatography with a Superdex 200 increase 10/300 column (GE Healthcare) in Tris buffered saline(TBS). Fab rich fractions were pooled and concentrated for EMPEM complex preparation.

### Preparation of Fab neuraminidase polyclonal complexes

A/Indiana/10/2011 N2 (60ug) was incubated with polyclonal fab (2mg) at room temperature overnight. The complex was SEC purified using a HiLoad 16/600 Superdex pg200 (GE Healthcare) column, with TBS as a running buffer. SEC fractions corresponding to the complex were pooled and concentrated to 0.5 mg/ml using an Amicon filter unit with 10 kDa cutoff (EMD Millipore)

### Preparation of grids for cryoEMPEM and imaging

UltrAuFoil R 1.2/1.3 grids (Au, 300 mesh; Quantifoil Micro ToolsGmbH) were used for sample vitrification. The grids were treated with Ar/O2 plasma (Solarus 950 plasma cleaner, Gatan) for 25s at 15mA immediately before sample application. A total of 0.5 μl of 0.7% w/v octyl-beta-glucoside(OBG) was added to 3.0 μl of the complex, and 3μl at 0.5mg/ml was immediately loaded onto the grid. Grids were prepared using Vitrobot mark IV (Thermo Fisher Scientific).

Temperature inside the chamber was maintained at 4°C, while humidity was at 100%. Blotting force was set to 1, wait time to 4.5s. Following the blotting step, the grids were plunge frozen into liquid ethane and cooled by liquid nitrogen. Grids were loaded into a Thermofisher Glacios microscope operating at 200kV. Movies were collected at 190,000x nominal magnification using a Falcon 4i camera using automated image collection software EPU 3.5.1 (ThermoFisher). Movies were aligned, dose-weighted and Contrast Transfer Function (CTF)-corrected in the CryoSPARC Live software platform.

### CryoEMPEM data processing

Data processing was carried out in CryoSPARC v4.5.1. Particles were picked using Template picker with a 200Å particle diameter using templates generated from class averages from CryoSPARC Live 2D classification jobs(*47*). Clean particles stacks were selected from further reference-free 2D classification jobs. These clean particles stacks were down sampled from their initial 512px box size to 64px using a downsample particles job and underwent Ab Initio reconstruction requesting 10 classes with a maximum alignment resolution of 20Å. Four immune complex volumes and one junk volume were selected for a subsequent heterogenous refinement with initial alignment resolution 25Å and maximum alignment resolution 20Å. Particles from the classes with active side binding antibodies were re-extracted at a box size of 512px and underwent non-uniform refinement(*48*) followed by local refinement using a NA-fv mask generated from a Volume tools job using an input volume generated in Chimera(*49*) with molmap and mask parameters: 0.05 map threshold, 3px dilation radius and 6px soft padding width. This final map was used in ModelAngelo analysis.

### CryoEMPEM model building with ModelAngelo

For HIV-1 benchmarking previously published maps were used. We used the maps deposited in the Electron Microscopy Databank (EMDB) as follows: for pAbC-1 EMD-23227, and for pAbC-2, EMD-23232. For the NA cryoEMPEM reported here map has been deposited in EMDB with ID EMD-48118. For all experiments reported here modelangelo v1.0.1 was used. All maps were run using the “model_angelo build_no_seq” command with default parameters. In many cases the chains and .hmm profiles that were output by MA were fragmented. In these cases, a poly-A fv model was aligned with the output.cif file in Pymol(*50*) to determine which .hmm profiles could be merged together to form a complete heavy or light chain .hmm file. To merge .hmm files generated by MA we generated auxiliary code that generates an amino acid probability array from the individual .hmm files from the fragments that can be merged into one complete array.

Then using the framework of the MA code, aaprobstohmm.py(*51*), that is responsible for generating .hmm files from the amino acid logits generated in an MA build_no_seq run, a new.hmm of the merged fragments can be generated. Where it was deemed that an amino acid was missing between fragments, we inserted a random amino acid into the .hmm file (5% probably of each of the 20 amino acids). A list of fragments that were pieced together can be found in this paper’s github repository. The output model’s .hmm profiles were used to search the unpaired BCR sequence database using HMMER(*52*).

### MAb selection using Model Angelo’s HMMER functionality

For HIV-1 benchmarking data where BCR-sequencing data was unpaired, the heavy and light sequences with the lowest E-score were selected for mAb synthesis. For the NA vaccination study, the heavy chains sequences with lowest E-scores for the top 4 predicted heavy V gene usages shown in Figure 2B were selected and for the light chain, three IGLV6-23 sequences with the highest E-scores and unique IGHV gene pairs were selected for mAb synthesis. Upon inspection of the top hits for IGHV5-6 and IGHV5-17 built into the cryoEMPEM map, we decided that the second lowest E-scoring sequences for these Vgenes fit better and thus moved forward with these mAbs for validation.

### MAb Expression

All monoclonal antibodies derived from MA analysis as well as the SFS mAbs previously described from the original SFS study were expressed using Genscript’s TurboCHO v2.0 service using 10ml expressions. Macaque fv sequences were synthesized into a human IgG1 backbone while mice fv sequences were synthesized into a mouse IgG1 backbone.

### Biolayer interferometry

IgG-binding kinetics were measured using biolayer interferometry on an Octet 96 Red instrument (ForteBio) using AHCII biosensors (Sartorius). IgGs were diluted to 5 μg ml^−1^ in kinetics buffer (20mM HEPES pH7.5, 0.02% Tween 20, 0.1% BSA). Each analyte was serially diluted to the desired concentration in kinetics buffer with the following final concentrations, 1000mM, 500mM, 250 mM, 125 mM, 62.3 mM,32.3 mM,15.6 mM. IgGs were loaded onto the biosensor for 60s. After loading, biosensors were placed in kinetics buffer for 60 s for a baseline reading. Biosensors were then placed in analyte for 1500s in the association phase, followed by 300s in the dissociation phase in kinetics buffer. The background signal from an unloaded sensor that underwent the same cycle with an analyte well of 1000mM, was subtracted from the other 7 signals. Kinetic analyses were performed twice with independently prepared analyte dilution series. Curve fitting was performed using a 2:1 binding model using the Octet data analysis studio (v13.0.2.46). Final plots were prepared using matplotlib.

### Virus production

A/Indiana/10/2011 (Ind11) H3N2 virus was obtained from International Reagent Resource (catalog# FR-947) and injected into 10-day old embryonated specific pathogen free (SPF) chicken eggs and incubated for 48 hours at 37°C. At the end of the incubation period, the allantoic fluid from the eggs was harvested and clarified by centrifugation at 300xg. The clarified allantoic fluid was then aliquoted and stored at -80 °C.

### ELISAs with neuraminidase

96 well high binding plates (SpectraPlate-96HB) were coated with recombinant NAs diluted in PBS at 2ug/mL (100uL per well) overnight at 4°C. Plates were washed 3 times with phosphate buffere saline with 0.05% v/v Tween-20(PBST) in the BioStack Microplate Stacker system (BioTek) (same washing conditions were used for all the following steps) and then blocked (200uL per well) with non-fat milk (5% w/v in PBST) for 1 hour (all the incubations were performed at room temperature). Primary antibodies were serial diluted (starting at 20ug/mL and followed by 1:5 dilutions) in PBST and aliquoted in plates (100uL per well) after the plates were washed. After 1 hour, the plates were washed again, and incubated with goat anti-mouse (Bio-Rad #STAR120P) or anti-human (Invitrogen #SA5-10283) IgG Fc-Horse Radish Peroxidase (HRP) secondary antibodies (1:20,000 diluted in PBST, 100uL per well) for 1 hour. Following PBST washing, 3,3’,5,5’-Tetramethylbenzidine (TMB) (Sigma #CLS2592) 2-Component Microwell Peroxidase Substrate Kit (Seracare #5120-0047) was used for color development according to manufacturer’s protocol. Synergy H1 plate reader (BioTek) was used for acquisition of colorimetric data by recording the absorbance at 450-nm wavelength. Data were analyzed with GraphPad Prism (version 10), and midpoint titers (EC_50_) were determined. All experiments were performed in duplicates.

### NA enzyme linked lectin assay (ELLA) with recombinant NA and Ind 11 H3N2 virus

96 well high binding plates (SpectraPlate-96HB) were coated with 25 ug/mL fetuin (Sigma #3385, 100 uL per well) diluted in coating buffer (KPL #50-84-01). To determine the optimal recombinant NA or Ind11 H3N2 virus concentrations used for ELLA, 50 uL of recombinant NA, serially diluted in Sample Diluent (PBS with CaCl_2_ & MgCl_2_ + 1% BSA + 0.5% Tween 20), starting from 20 ug/mL followed by 1:3 dilutions, or 150 uL of 1:2 serially diluted A/Indiana/10/2011 (Ind 11) H3N2 virus was transferred to fetuin-coated plates after washing 3 times with PBST. The plates were then sealed and incubated at 37 °C for 16-18 hours with the recombinant NA or were incubated at 33 °C for 6 hours with the A/Indiana/10/2011 (H3N2) virus. Next, plates were washed 5 times with PBST before adding 100 uL/well of 1 ug/mL peanut agglutinin (PNA)-HRP (Sigma #7759-1mg, diluted in Sample Diluent w/o Tween 20) in each well. The plates were incubated at room temperature for 2 hours and then washed 3 times with PBST. The color was developed with the same TMB substrate kit used in ELISA above. Data were analyzed with GraphPad Prism (version 10), and a standard curve was determined. The NA concentration within the linear region of the standard curve that conferred the OD450 closest to the max signal was selected for ELLA (the selected signal should also be at least 10- fold higher than background signal). To determine the NA activity inhibition of the antibodies, 25 uL serial diluted antibodies (1:2 dilution starting from 20 ug/mL or 30 ug/mL) were mixed with 25 uL NA (final NA concentration should match that was determined above). The 50 uL mixture was then transferred to each well in washed fetuin-coated plates. The rest of the ELLA assay was finished by following the same steps described above. The NA activity inhibition was quantified as: 100%-(signal of sample well/signal of NA-only well) × 100%, with the NA-only well has no inhibition (0%). Data were analyzed with GraphPad Prism (version 10), and midpoint inhibitory titers (IC_50_) were determined. All experiments were performed in duplicates.

### Mice protection experiment

Six-to-eight-week-old female DBA/2J mice from Jackson Laboratories were used for all animal experiments. All procedures were conducted in accordance with protocols approved by the Institutional Animal Care and Use Committee (IACUC) at the Icahn School of Medicine at Mount Sinai.

For the experiment, six groups of three to five mice each were administered intraperitoneally with 5 mg/mL of generated anti-NA mAbs; 5-12, 5-16, 6-23.1, and 6-23.2, a well-defined anti-NA antibody (1G01) was used as a positive control, and an irrelevant antibody (anti-H5) was used as a negative control. Two hours after the mAb administration, mice were intranasally infected with 5 times the 50% lethal dose (LD_50_) of the Ind 11 H3N2 virus under anesthesia, and weight loss was monitored for 14 days post-infection. Mice that lost 25% or more of their initial body weight were humanely euthanized. The sample size for the indicated groups were: 5-12(n=4), 5-16(n=5), 6-23.1(n=3), 6.23-2(n=5), 1G01(n=5), and anti-H5(n=5).

### CryoEM analysis of mAb complexes:Preparation of mAb NA complex grids

Ind11 NA was incubated overnight at 4°C with 2x molar excess of mAb IgG per protomer at a final concentration of 0.6mg/ml in TBS buffer. 1.2/1.3 copper Quantifoil 300 mesh grid (Quantifoil Micro ToolsGmbH) were used for sample vitrification. The grids were treated with Ar/O2 plasma (Solarus 950 plasma cleaner, Gatan) for 30s at 15mA immediately before sample application. 0.5ul of 0.7% w/v OBG detergent was added to 3ul of the complex, and 3 ul was immediately loaded onto the grid. Grids were prepared using Vitrobot mark IV (Thermo Fisher Scientific).Temperature inside the chamber was maintained at 4°C, while humidity was at 100%. Blotting force was set to 1, wait time to 5s, while the blotting time was varied within a 3.5-5s range. Following the blotting step, the grids were plunge frozen into liquid ethane and cooled by liquid nitrogen.

### CryoEM analysis of mAb complexes:Data Collection and Processing

Complexes were imaged at 190,000× nominal magnification using a Falcon 4i camera on a Glacios microscope at 200 kV. Automated image collection was performed using EPU 3.5.1 (ThermoFisher). Images were aligned, dose-weighted, and Contrast Transfer Function (CTF)-corrected in the CryoSPARC Live™ software platform. Briefly, particles were picked using templates generated from the live 2D classification job and extracted at 512px box size, fourier cropped to 256px. Clean particle stacks were selected through reference-free 2D classification, which were then used to generate a 3D reference model from *ab initio* job, which was then initially refined using either Non-Uniform refinement or homogenous refinement with C4 symmetry. Final maps were generated with C4 symmetry using local refinement in CryoSparc 4.5.1. The number of initial and final particles is mentioned in Supplementary Figure 5.

### Model Building and refinement

An unpublished Ind11 NA model was used as an initial model for NA and mAb models were generated using Abodybuilder-2(*29*). The initial models were first rigid body fitted into the cryo-EM maps in ChimeraX(*53*) and the fitting was validated in COOT 0.9.8.7(*30*). The models were then refined using real-space refinement in PHENIX(*54*) and COOT iteratively. The model refinement statistics are shown in Supplementary Figure 5. The Kabat numbering system was used for the mAbs, and the N2 numbering convention is used for the alignment of NA.

## Acknowledgments

The authors thank Mr. Will Lessen, Dr. Anant Ghapure, and Ms. Hannah Turner from the Scripps Research Institute for their help with EM experiments. We thank Dr. Lauren Holden for their help in preparing this manuscript. We thank Dr. Gabe Ozorowski and Dr. Aleks Antanasijevic for helpful conversations regarding HIV-1 and the structure from sequence method. We thank Dr. Jeffery Copps and Ms. Grace Gibson for HIV-1 Envelope protein, and Mr. Charles Bowman and Mr. JC Ducom for compute support. Work in the Krammer laboratory was partially funded by the Centers of Excellence for Influenza Research and Response (CEIRR, contract # 75N93021C00014, reagent generation) and by institutional funds.

## Data Availability

The data that support this study are available from the corresponding authors upon request. The cryo-EM maps and fitted coordinates have been deposited to the Electron Microscopy Data Bank (EMDB) and Protein Data Bank (PDB) with accession codes listed in Supplemental Figure 5.

## Code Availability

The hmm merger scripts will be open source upon publication and is also available upon request during the review process.

## References

1. www.antibodysociety.org/antibody-therapeutics-product-data.

2. M. S. Castelli, P. McGonigle, P. J. Hornby, The pharmacology and therapeutic applications of monoclonal antibodies. Pharmacol Res Perspect 7, e00535 (2019).

3. E. Lacorte, A. Ancidoni, V. Zaccaria, G. Remoli, L. Tariciotti, G. Bellomo, F. Sciancalepore, M. Corbo, F. L. Lombardo, I. Bacigalupo, M. Canevelli, P. Piscopo, N. Vanacore, Safety and Efficacy of Monoclonal Antibodies for Alzheimer’s Disease: A Systematic Review and Meta-Analysis of Published and Unpublished Clinical Trials. Journal of Alzheimer’s Disease 87, 101–129 (2022).

4. M. Kothari, A. Wanjari, S. Acharya, V. Karwa, R. Chavhan, S. Kumar, A. Kadu, R. Patil, A Comprehensive Review of Monoclonal Antibodies in Modern Medicine: Tracing the Evolution of a Revolutionary Therapeutic Approach. Cureus 16, e61983 (2024).

5. M. Ballow, R. Ortiz-de-Lejarazu, I. Quinti, M. S. Miller, K. Warnatz, Contribution of immunoglobulin products in influencing seasonal influenza infection and severity in antibody immune deficiency patients receiving immunoglobulin replacement therapy. Front Immunol 15, 1452106 (2024).

6. G. Köhler, C. Milstein, Continuous cultures of fused cells secreting antibody of predefined specificity. Nature 256, 495–497 (1975).

7. J. McCafferty, A. D. Griffiths, G. Winter, D. J. Chiswell, Phage antibodies: filamentous phage displaying antibody variable domains. Nature 348, 552–554 (1990).

8. A. Doerner, L. Rhiel, S. Zielonka, H. Kolmar, Therapeutic antibody engineering by high efficiency cell screening. FEBS Lett 588, 278–287 (2014).

9. J. Zhu, D. Hatton, New mammalian expression systems. Adv Biochem Eng Biotechnol 165, 9–50 (2018).

10. H. A. Utset, J. J. Guthmiller, P. C. Wilson, Bridging the B Cell Gap: Novel Technologies to Study Antigen-Specific Human B Cell Responses. Vaccines (Basel) 9 (2021).

11. A. Madsen, Y.-N. Dai, M. McMahon, A. J. Schmitz, J. S. Turner, J. Tan, T. Lei, W. B. Alsoussi, S. Strohmeier, M. Amor, B. M. Mohammed, P. A. Mudd, V. Simon, R. J. Cox, D. H. Fremont, F. Krammer, A. H. Ellebedy, Human antibodies targeting influenza B virus neuraminidase active site are broadly protective. Immunity 53, 852–863.e7 (2020).

12. K. M. McIntire, H. Meng, T. H. Lin, W. Kim, N. E. Moore, J. Han, M. McMahon, M. Wang, S. K. Malladi, B. M. Mohammed, J. Q. Zhou, A. J. Schmitz, K. B. Hoehn, J. M. Carreño, T. Yellin, T. Suessen, W. D. Middleton, S. A. Teefey, R. M. Presti, F. Krammer, J. S. Turner, A. B. Ward, I. A. Wilson, S. H. Kleinstein, A. H. Ellebedy, Maturation of germinal center B cells after influenza virus vaccination in humans. J Exp Med 221 (2024).

13. J. Wrammert, K. Smith, J. Miller, W. A. Langley, K. Kokko, C. Larsen, N. Y. Zheng, I. Mays, L. Garman, C. Helms, J. James, G. M. Air, J. D. Capra, R. Ahmed, P. C. Wilson, Rapid cloning of high-affinity human monoclonal antibodies against influenza virus. Nature 453, 667–671 (2008).

14. P. O. Byrne, J. S. McLellan, Principles and practical applications of structure-based vaccine design. Curr Opin Immunol 77, 102209 (2022).

15. S. Bangaru, A. Antanasijevic, N. Kose, L. M. Sewall, A. M. Jackson, N. Suryadevara, X. Zhan, J. L. Torres, J. Copps, A. T. de la Peña, J. E. Crowe, A. B. Ward, Structural mapping of antibody landscapes to human betacoronavirus spike proteins. Sci Adv 8, 2911 (2022).

16. A. Antanasijevic, L. M. Sewall, C. A. Cottrell, D. G. Carnathan, L. E. Jimenez, J. T. Ngo, J. B. Silverman, B. Groschel, E. Georgeson, J. Bhiman, R. Bastidas, C. LaBranche, J. D. Allen, J. Copps, H. R. Perrett, K. Rantalainen, F. Cannac, Y. R. Yang, A. T. de la Peña, R. F. Rocha, Z. T. Berndsen, D. Baker, N. P. King, R. W. Sanders, J. P. Moore, S. Crotty, M. Crispin, D. C. Montefiori, D. R. Burton, W. R. Schief, G. Silvestri, A. B. Ward, Polyclonal antibody responses to HIV Env immunogens resolved using cryoEM. Nat Commun 12 (2021).

17. R. Ray, F. A. Nait Mohamed, D. P. Maurer, J. Huang, B. A. Alpay, L. Ronsard, Z. Xie, J. Han, M. Fernandez-Quintero, Q. A. Phan, R. L. Ursin, M. Vu, K. H. Kirsch, T. Prum, V. C. Rosado, T. Bracamonte-Moreno, V. Okonkwo, J. Bals, C. McCarthy, U. Nair, M. Kanekiyo, A. B. Ward, A. G. Schmidt, F. D. Batista, D. Lingwood, Eliciting a single amino acid change by vaccination generates antibody protection against group 1 and group 2 influenza A viruses. Immunity 57, 1141–1159.e11 (2024).

18. A. Antanasijevic, C. A. Bowman, R. N. Kirchdoerfer, C. A. Cottrell, G. Ozorowski, A. A. Upadhyay, K. M. Cirelli, D. G. Carnathan, C. A. Enemuo, L. M. Sewall, B. Nogal, F. Zhao, B. Groschel, W. R. Schief, D. Sok, G. Silvestri, S. Crotty, S. E. Bosinger, A. B. Ward, From structure to sequence: Antibody discovery using cryoEM. Sci Adv 8, 2039 (2022).

19. K. Jamali, L. Käll, R. Zhang, A. Brown, D. Kimanius, S. H. W. Scheres, Automated model building and protein identification in cryo-EM maps. Nature *2024 628:8007* 628, 450–457 (2024).

20. S. R. Eddy, Accelerated Profile HMM Searches. PLoS Comput Biol 7, e1002195 (2011).

21. R. W. Sanders, M. Vesanen, N. Schuelke, A. Master, L. Schiffner, R. Kalyanaraman, M. Paluch, B. Berkhout, P. J. Maddon, W. C. Olson, M. Lu, J. P. Moore, Stabilization of the soluble, cleaved, trimeric form of the envelope glycoprotein complex of human immunodeficiency virus type 1. J Virol 76, 8875–8889 (2002).

22. T. J. Wheeler, J. Clements, R. D. Finn, Skylign: A tool for creating informative, interactive logos representing sequence alignments and profile hidden Markov models. BMC Bioinformatics 15, 1–9 (2014).

23. https://www.imgt.org.

24. T. H. Olsen, I. H. Moal, C. M. Deane, Addressing the antibody germline bias and its effect on language models for improved antibody design. bioRxiv,.02.02.578678 (2024).

25. B. Abanades, W. K. Wong, F. Boyles, G. Georges, A. Bujotzek, C. M. Deane, ImmuneBuilder: Deep-Learning models for predicting the structures of immune proteins. Communications Biology *2023 6:1* 6, 1–8 (2023).

26. P. Emsley, M. Crispin, Structural analysis of glycoproteins: Building N-linked glycans with coot. Acta Crystallogr D Struct Biol 74, 256–263 (2018).

27. https://getcontacts.github.io/.

28. P. J. A. Cock, T. Antao, J. T. Chang, B. A. Chapman, C. J. Cox, A. Dalke, I. Friedberg, T. Hamelryck, F. Kauff, B. Wilczynski, M. J. L. De Hoon, Biopython: freely available Python tools for computational molecular biology and bioinformatics. Bioinformatics 25, 1422–1423 (2009).

29. P. J. M. Brouwer, H. R. Perrett, T. Beaumont, H. Nijhuis, S. Kruijer, J. A. Burger, W.-H. Lee, H. Müller-Kraüter, R. W. Sanders, T. Strecker, M. J. van Gils, A. B. Ward, Defining bottlenecks and opportunities for Lassa virus neutralization by structural profiling of vaccine-induced polyclonal antibody responses. bioRxiv, 2023.12.21.572918 (2023).

30. S. Brown, A. Antanasijevic, L. M. Sewall, D. M. Garcia, P. J. M. Brouwer, R. W. Sanders, A. B. Ward, Anti-Immune Complex Antibodies are Elicited During Repeated Immunization with HIV Env Immunogens. bioRxiv, 2024.03.15.585257 (2024).

31. D. Schulte, M. Šiborová, L. Käll, J. Snijder, Simultaneous polyclonal antibody sequencing and epitope mapping by cryo electron microscopy and mass spectrometry – a perspective. bioRxiv, 2024.06.21.600107 (2024).

32. M. Haraldson Høie, A. M. Hummer, T. H. Olsen, M. Nielsen, C. M. Deane, AntiFold: Improved antibody structure design using inverse folding. https://opig.stats.ox.ac.uk/data/downloads/AntiFold.

33. J. Dauparas, I. Anishchenko, N. Bennett, H. Bai, R. J. Ragotte, L. F. Milles, B. I. M. Wicky, A. Courbet, R. J. de Haas, N. Bethel, P. J. Y. Leung, T. F. Huddy, S. Pellock, D. Tischer, F. Chan, B. Koepnick, H. Nguyen, A. Kang, B. Sankaran, A. K. Bera, N. P. King, D. Baker, Robust deep learning–based protein sequence design using ProteinMPNN. Science *(1979)* 378, 49–56 (2022).

34. F. A. Dreyer, D. Cutting, C. Schneider, H. Kenlay, C. M. Deane, Inverse folding for antibody sequence design using deep learning. (2023).

35. D. Stadlbauer, X. Zhu, M. McMahon, J. S. Turner, T. J. Wohlbold, A. J. Schmitz, S. Strohmeier, W. Yu, R. Nachbagauer, P. A. Mudd, I. A. Wilson, A. H. Ellebedy, F. Krammer, Broadly protective human antibodies that target the active site of influenza virus neuraminidase. Science 366, 499 (2019).

36. J. J. Lavinder, A. P. Horton, G. Georgiou, G. C. Ippolito, Next-generation sequencing and protein mass spectrometry for the comprehensive analysis of human cellular and serum antibody repertoires. Curr Opin Chem Biol 24, 112–120 (2015).

37. I. Snapkov, M. Chernigovskaya, P. Sinitcyn, K. Lê Quý, T. A. Nyman, V. Greiff, Progress and challenges in mass spectrometry-based analysis of antibody repertoires. Trends Biotechnol 40, 463–481 (2022).

38. L. Wossnig, N. Furtmann, A. Buchanan, S. Kumar, V. Greiff, Best practices for machine learning in antibody discovery and development. Drug Discov Today 29, 104025 (2024).

39. G. Jo, S. Yamayoshi, K. M. Ma, O. Swanson, J. L. Torres, J. A. Ferguson, M. L. Fernandez-Quintero, J. Huang, J. Copps, A. J. Rodriguez, J. M. Steichen, Y. Kawaoka, J. Han, A. B. Ward, Structural basis of broad protection against influenza virus by a human antibody targeting the neuraminidase active site via a recurring motif in CDR H3. bioRxiv, 2024.11.26.625467 (2024).

40. Z. Xie, Y. C. Lin, J. M. Steichen, G. Ozorowski, S. Kratochvil, R. Ray, J. L. Torres, A. Liguori, O. Kalyuzhniy, X. Wang, J. E. Warner, S. R. Weldon, G. A. Dale, K. H. Kirsch, U. Nair, S. Baboo, E. Georgeson, Y. Adachi, M. Kubitz, A. M. Jackson, S. T. Richey, R. M. Volk, J. H. Lee, J. K. Diedrich, T. Prum, S. Falcone, S. Himansu, A. Carfi, J. R. Yates, J. C. Paulson, D. Sok, A. B. Ward, W. R. Schief, F. D. Batista, mRNA-LNP HIV-1 trimer boosters elicit precursors to broad neutralizing antibodies. Science *(1979)* 384 (2024).

41. Y. Hao, S. Hao, E. Andersen-Nissen, W. M. Mauck, S. Zheng, A. Butler, M. J. Lee, A. J. Wilk, C. Darby, M. Zager, P. Hoffman, M. Stoeckius, E. Papalexi, E. P. Mimitou, J. Jain, A. Srivastava, T. Stuart, L. M. Fleming, B. Yeung, A. J. Rogers, J. M. McElrath, C. A. Blish, R. Gottardo, P. Smibert, R. Satija, Integrated analysis of multimodal single-cell data. Cell 184, 3573–3587.e29 (2021).

42. H. L. Turner, A. M. Jackson, S. T. Richey, L. M. Sewall, A. Antanasijevic, L. Hangartner, A. B. Ward, Protocol for analyzing antibody responses to glycoprotein antigens using electron-microscopy-based polyclonal epitope mapping. STAR Protoc 4, 102476 (2023).

43. A. Punjani, J. L. Rubinstein, D. J. Fleet, M. A. Brubaker, cryoSPARC: algorithms for rapid unsupervised cryo-EM structure determination. Nature Methods *2017 14:3* 14, 290–296 (2017).

44. A. Punjani, H. Zhang, D. J. Fleet, Non-uniform refinement: adaptive regularization improves single-particle cryo-EM reconstruction. Nature Methods *2020 17:12* 17, 1214–1221 (2020).

45. E. F. Pettersen, T. D. Goddard, C. C. Huang, G. S. Couch, D. M. Greenblatt, E. C. Meng, T. E. Ferrin, UCSF Chimera--a visualization system for exploratory research and analysis. J Comput Chem 25, 1605–1612 (2004).

46. The PyMOL Molecular Graphics System, Version 3.0 Schrödinger, LLC.

47. https://github.com/3dem/model-angelo/blob/main/model_angelo/utils/aa_probs_to_hmm.py.

48. R. D. Finn, J. Clements, W. Arndt, B. L. Miller, T. J. Wheeler, F. Schreiber, A. Bateman, S. R. Eddy, HMMER web server: 2015 update. Nucleic Acids Res 43, W30–W38 (2015).

49. E. C. Meng, T. D. Goddard, E. F. Pettersen, G. S. Couch, Z. J. Pearson, J. H. Morris, T. E. Ferrin, UCSF ChimeraX: Tools for structure building and analysis. Protein Science 32, e4792 (2023).

50. D. Liebschner, P. V. Afonine, M. L. Baker, G. Bunkoczi, V. B. Chen, T. I. Croll, B. Hintze, L. W. Hung, S. Jain, A. J. McCoy, N. W. Moriarty, R. D. Oeffner, B. K. Poon, M. G. Prisant, R. J. Read, J. S. Richardson, D. C. Richardson, M. D. Sammito, O. V. Sobolev, D. H. Stockwell, T. C. Terwilliger, A. G. Urzhumtsev, L. L. Videau, C. J. Williams, P. D. Adams, Macromolecular structure determination using X-rays, neutrons and electrons: recent developments in Phenix. Acta Crystallogr D Struct Biol 75, 861–877 (2019).

