## Supplemental Material for "Functional and epitope specific monoclonal antibody discovery directly from immune sera using cryoEM"

| pAb name | CryoEM map | MA no seq model | Chains used for HMMER search |
|---|---|---|---|
| **pAbC-1** | 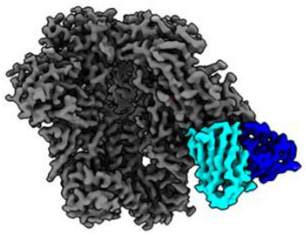 | 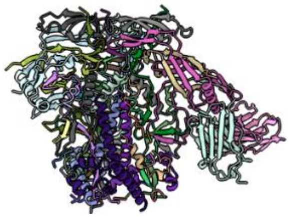 | 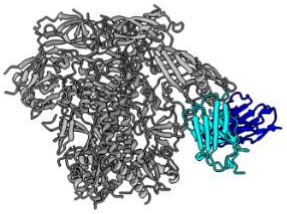 |
| **pAbC-2** | 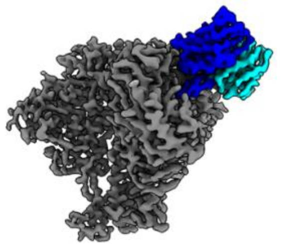 | 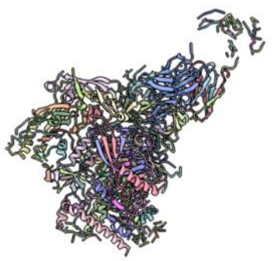 | 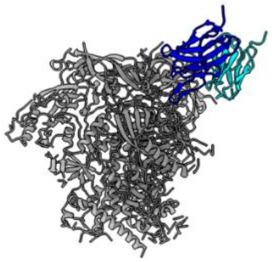 |

Supplemental Figure S1. Results of MA build_no_seq for pAbC-1 and pAbC-2. The cryoEM map column shows the input map used with the antigen regions shown in gray and regions corresponding to the heavy and light chain shown in dark blue and cyan, respectively. MA no seq model columns shows the output.cif generated by MA. For pAbC-2 chains corresponding to heavy and light chain were fragmented and were traced by docking in a polyA fv model. The last column shows the final chains, with corresponding .hmm files, used in the HMMER search (heavy and light are in blue and cyan respectively).

```
PABC-1_ORIGINAL_HC        QVQLQESGPGLVKPSETLSLTCAVSGGSFSGYSWGWIRQPPGKGLEWIGSIIGRTGSTAY   60
PABC-1_MA_OUTPUT_HC       SVQIQITGEDLVKPDKTLTVTCSVEGGGHSGYSYGYVRQPPGKGLEYIGMIIGRSGETDY   60
                          .**:* :* .****.:**::**:*.**..****:*::*********:** ****:*.* *

PABC-1_ORIGINAL_HC        NPSLTSRVTISRDTSNNQFSLKLTSLTAADTAVYYCARQQSNFDFWGQGVLVTVSS-     116
PABC-1_MA_OUTPUT_HC       NPKLEPRVTISRDESKNQYSLKLTDVTSGDNGVYYCGRQQAIFDYWGQGLLVTRSDV     117
                          **.*  ******* *:**:*****.:*:.*..****.***: **:****:*** *.
```

```
PABC-1_ORIGINAL_LC        DIQMTQSPSSLSASVGDTVTTTCRASQDISNDLAWYQQKPGKAPKPLLYYASNLESGVPS   60
PABC-1_MA_OUTPUT_LC       AINMTQSPTTLSGEKGETVTCTCRATQEITNDLAWFQQKPGKSPKPLIYRASNLEVGVPS   60
                           *:*****::**.. *:*** ****:*:*:*****:******:****:* ***** ****

PABC-1_ORIGINAL_LC        MFSGSGSGTDFTLTISSLQPEDFASYFCQQYNSYPRTFGQGTKVEIK       107
PABC-1_MA_OUTPUT_LC       RYSGTGSGDEFTLTISDLNPESRAIYYCMNHHERPWVFGPGTKISNR       107
                           :**:*** :******.*:**. * *:* :::. * .** ***:. :
```

```
PABC-2_ORIGINAL_HC        QVQLVQSGAEVKMPGTSVKLSCKTSGYTFTSYNINWVRQAPG-QALEWMGWINPNNGTTD   59
PABC-2_MA_OUTPUT_HC       NRQLTQAGSAVKKPGESVKLSCKAAGRNFSAYNINWVRQADGKQALEWMGYLNPENGQEE   60
                          : **.*:*: ** ** *******::* .*::********* * *******::**:**  :

PABC-2_ORIGINAL_HC        YAQKFQGRVTMTRDTSTTTAYMQLNSLRSEDTAVYYCARARGGYEDDDGYHYTGYGLDSW   119
PABC-2_MA_OUTPUT_HC       YSEEFEGRVTFSRDTETNEVYLQLKNLKVENTSIYYCARARAGYEDEEGFHYTGYGMDTF   120
                          *:::*:****::***.*. .*:**:.*: *:*::*******.****::*:******:*::

PABC-2_ORIGINAL_HC        GQGVVVTVSS-    129
PABC-2_MA_OUTPUT_HC       GQAVVIEVSPS    131
                          **.**: **
```

```
PABC-2_ORIGINAL_LC        --DIQMTQSPSSLSASIGDRVTVTCRASQGINMQLCWYQLKPGKAPTLLIYGTSGLQTGV   58
PABC-2_MA_OUTPUT_LC       GEKIKMTQSPSXXSSSLGDRVTVTCRAAEGNENELSWYKQLPGKPPTLLIYGADGINSGV   60
                            .*:******  *:*:**********::* : :*.**:  *** *******:.*:::**

PABC-2_ORIGINAL_LC        SSRFSGSGSGTNFTLTISSLQ-PEDVATYYCQQDYTTPFTFGPGTKLDIK--   107
PABC-2_MA_OUTPUT_LC       SPRFSGSGGDNDFSLTLSSLNNPNDIGVFYQMQCHSPVFTFGPGVEVPEGLS   112
                          * ******...:*:**:***: *:*:..:*  * ::  ******.::
```

Supplemental Figure S2. Sequence identity comparisons of MA build_no_seq model output vs mAbs originally identified from manual structure to sequence(SFS) paper.

```
PABC-1_ORIGINAL_HC      QVQLQESGPGLVKPSETLSLTCAVSGGSFSGYSWGWIRQPPGKGLEWIGSIIGRTGSTAY    60
PABC-1_TOPHIT_HC        QVQLQESGPGLVKPSETLSLTCAVSGGSFSGYSWGWIRQPPGKGLEWIGYIIGRTGSTDY    60
                        ************************************************* ******** *

PABC-1_ORIGINAL_HC      NPSLTSRVTISRDTSNNQFSLKLTSLTAADTAVYYCARQQSNFDFWGQGVLVTVSS        116
PABC-1_TOPHIT_HC        NPSLTSRVTISRDTSNNQFSLKLTSLTAADTAVYYCARQQSNFDFWGQGVLVTVSS        116
                        ********************************************************
```

```
PABC-1_ORIGINAL_LC      DIQMTQSPSSLSASVGDTVTTTCRASQDISNDLAWYQQKPGKAPKPLLYYASNLESGVPS    60
PABC-1_TOPHIT_LC        DIQMTQSPSSLSASVGDTVTATCRASQDITNDLAWYQQKPGKAPKPLIYYASNLESGVPS    60
                        ********************:********:*****************:************

PABC-1_ORIGINAL_LC      MFSGSGSGTDFTLTISSLQPEDFASYFCQQYNSYPRTFGQGTKVEIK 107
PABC-1_TOPHIT_LC        RFSGSGSGTDFTLTISSLQPEDFAIYFCQQFYTYPRTFGQGTKVEIK 107
                         *********************** *****: :**************
```

```
PABC-2_ORIGINAL_HC      QVQLVQSGAEVKMPGTSVKLSCKTSGYTFTSYNINWVRQAPGQALEWMGWINPNNGTTDY    60
PABC-2_TOPHIT_HC        QVQLVQSGAEVKMPGTSVKLSCKTSGYTFTSYNINWVRQAPGQALEWMGWINPKNGKTDY    60
                        *****************************************************:**.***

PABC-2_ORIGINAL_HC      AQKFQGRVTMTRDTSTTTAYMQLNSLRSEDTAVYYCARARGGYEDDDGYHYTGYGLDSWG    120
PABC-2_TOPHIT_HC        AKKFQGRVTMTRDTSTTTAYMQLNSLRSEDTAVYYCARARGGYEDDDGYHYTGYGLDSWG    120
                        *:**********************************************************

PABC-2_ORIGINAL_HC      QGVVVTVSS       129
PABC-2_TOPHIT_HC        QGVVVTVSS       129
                        *********
```

```
PABC-2_ORIGINAL_LC      DIQMTQSPSSLSASIGDRVTVTCRASQGINMQLCWYQLKPGKAPTLLIYGTSGLQTGVSS    60
PABC-2_TOPHIT_LC        DIQMTQSPSSLSASIGDRVTVTCRASQGIKKELSWFQQRPGRAPTLLIYGASSLQTGVST    60
                        *****************************: :*.*:* :**:********:*.******:

PABC-2_ORIGINAL_LC      RFSGSGSGTNFTLTISSLQPEDVATYYCQQDYTTPFTFGPGTKLDIK 107
PABC-2_TOPHIT_LC        RFSGSGSGTDFTLTISSLQPEDVATYYCQQDFSPPFTFGVGTKVEIK 107
                        *********:*********************:: ***** ***::**
```

Supplemental Figure S3. Sequence identity comparisons of ModelAngelo top hits from HMMER vs mAbs originally identified from manual SFS paper.

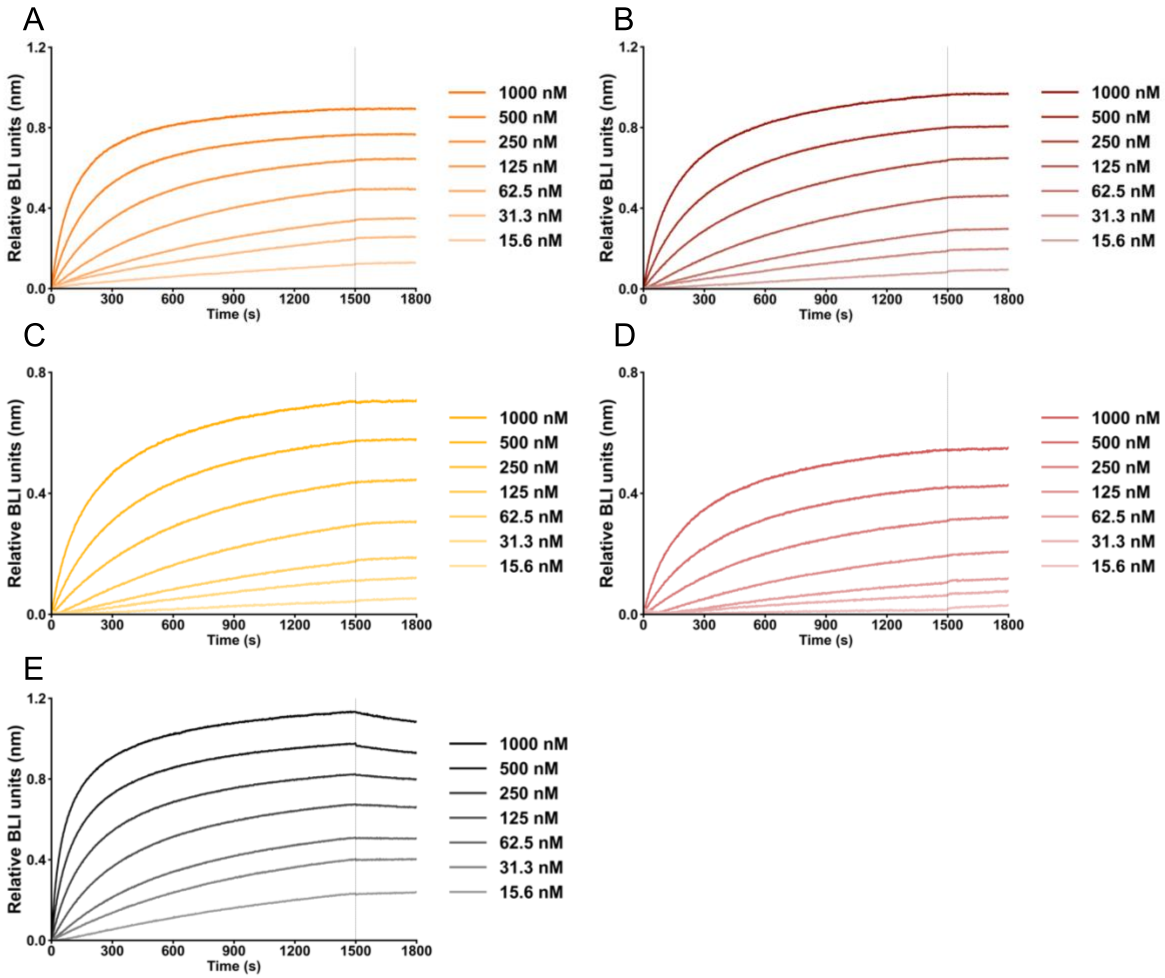


Supplemental Figure S4. Complete dilution series of BLI binding curves for determination of kinetic binding parameters of interactions between BG505 SOSIP (A) pAbC-1_ModelAngelo (B) pAbC-1_Manually derived (C) pAbC-2_ModelAngelo (D) pAbC-2_Manually derived (E) PGT-145.

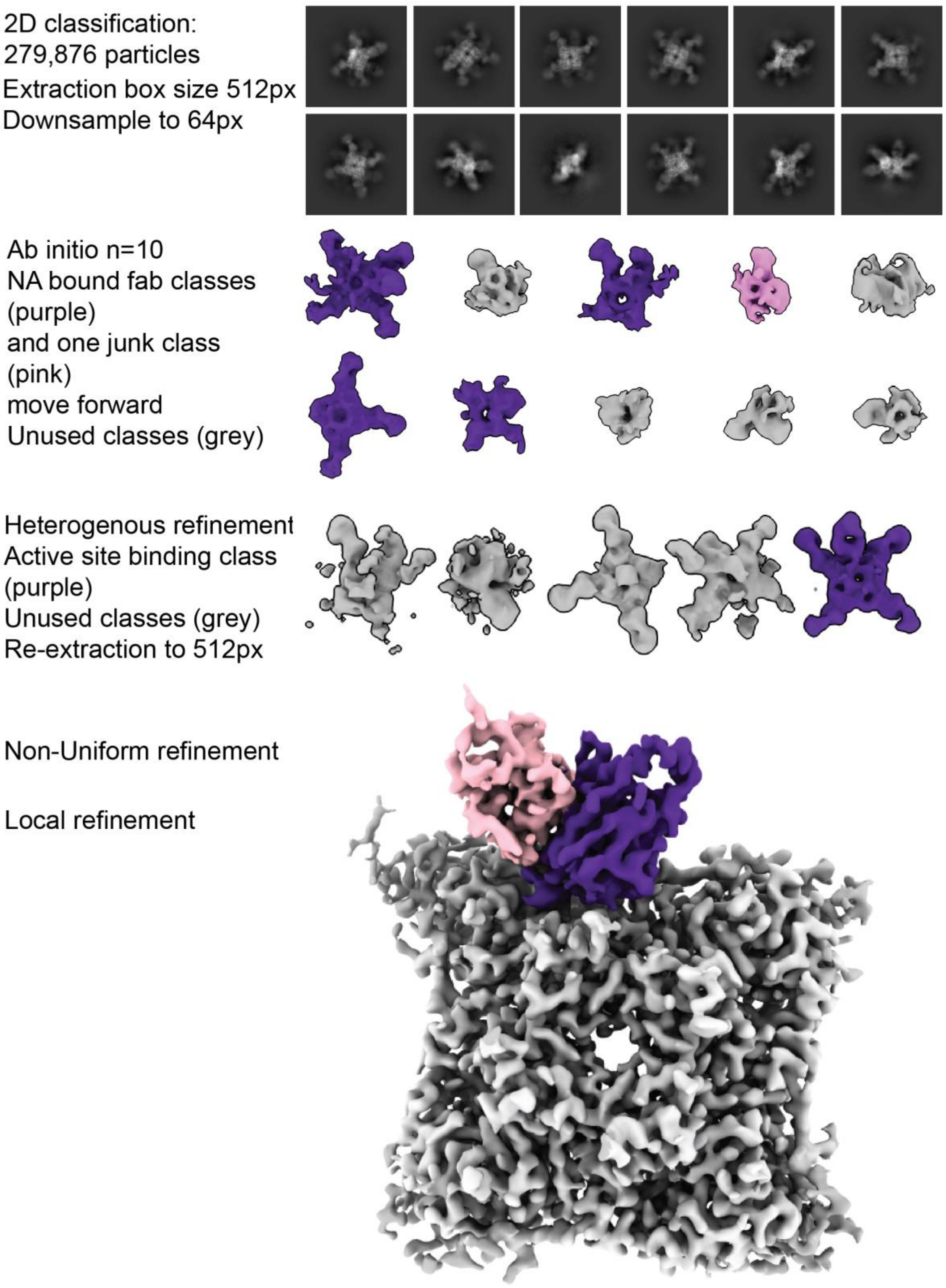


Supplemental Figure S5. cryoEMPEM processing workflow in cryosparc v4.3.0

| | NA in complex with polyclonal antibody (EMDB-48118) | NA in complex with 5-6 (EMDB-48165) (PDB-9MD2) | NA in complex with 5-12 (EMDB-48166) (PDB-9MD3) | NA in complex with 5-16 (EMDB-48167) (PDB-9MD4) | NA in complex with 6-23.2 (EMDB-48168) (PDB-9MD5) | NA in complex with 6-23.1 (EMDB-48169) (PDB-9MD6) |
|---|---|---|---|---|---|---|
| **Data collection and processing** | | | | | | |
| Magnification | 190000 | 190000 | 190000 | 190000 | 190000 | 190000 |
| Voltage (kV) | 200 | 200 | 200 | 200 | 200 | 200 |
| Electron exposure (e–/Å$^2$) | 43.4 | 45 | 45 | 45.1 | 45.12 | 40.9 |
| Defocus range (μm) | -1.8 to -0.8 | -1.8 to -0.8 | -1.8 to -0.8 | -1.8 to -0.8 | -1.8 to -0.8 | -1.8 to -0.8 |
| Pixel size (Å) | 0.725 | 0.725 | 0.725 | 0.725 | 0.725 | 0.718 |
| Symmetry imposed | C1 | C4 | C4 | C4 | C4 | C4 |
| Initial particle images (no.) | 436610 | 593280 | 740197 | 474585 | 254209 | 561753 |
| Final particle images (no.) | 72780 | 79388 | 67241 | 117508 | 77151 | 91670 |
| Map resolution (Å) | 3.3 | 3.4 | 2.9 | 2.7 | 2.9 | 2.7 |
| FSC threshold (0.143) | | | | | | |
| **Refinement** | | | | | | |
| Model resolution (Å) | - | 3.4 | 3.3 | 2.8 | 3.0 | 2.8 |
| FSC threshold (0.5) | | | | | | |
| Map sharpening *B* factor (Å$^2$) | - | -115.8 | -71.6 | -91.3 | -74.5 | -76.3 |
| Map Correlation Coefficient (Mask) | - | 0.85 | 0.87 | 0.89 | 0.87 | 0.89 |
| Model composition | | | | | | |
| Non-hydrogen atoms | - | 19540 | 20020 | 19688 | 19884 | 19532 |
| Protein residues | - | 2468 | 2468 | 2472 | 2472 | 2468 |
| Ligands | - | CA:8 | CA:4 | CA:4 | CA:4 | CA:4 |
| | - | NAG:24 | NAG:40 | NAG:24 | NAG:32 | NAG:20 |
| | - | BMA:4 | BMA:8 | BMA:4 | BMA:4 | BMA:4 |
| | | MAN:8 | MAN:20 | MAN:8 | MAN:8 | MAN:8 |
| *B* factors (Å$^2$) | | | | | | |
| Protein | - | 18.66 | 19.50 | 28.41 | 22.51 | 18.57 |
| Ligand | - | 13.91 | 23.04 | 28.04 | 23.02 | 32.83 |
| R.m.s. deviations | | | | | | |
| Bond lengths (Å) | - | 0.004 | 0.002 | 0.003 | 0.006 | 0.002 |
| Bond angles (°) | - | 0.968 | 0.578 | 0.618 | 0.756 | 0.560 |
| Validation | | | | | | |
| MolProbity score | - | 1.45 | 1.45 | 1.43 | 1.74 | 1.3 |
| Clashscore | - | 8.02 | 7.89 | 7.86 | 9.12 | 5.6 |
| Poor rotamers (%) | - | 0 | 0 | 0 | 0 | 0 |
| Cβ outliers | - | 0 | 0 | 0 | 0 | 0 |
| Ramachandran plot | | | | | | |
| Favored (%) | - | 97.95 | 97.91 | 98.20 | 96.2 | 98.69 |
| Allowed (%) | - | 1.88 | 1.92 | 1.47 | 3.64 | 1.15 |
| Disallowed (%) | - | 0.16 | 0.16 | 0.33 | 0.16 | 0.16 |

Supplemental Figure S6. Table of statistics for pAb map and mAbs

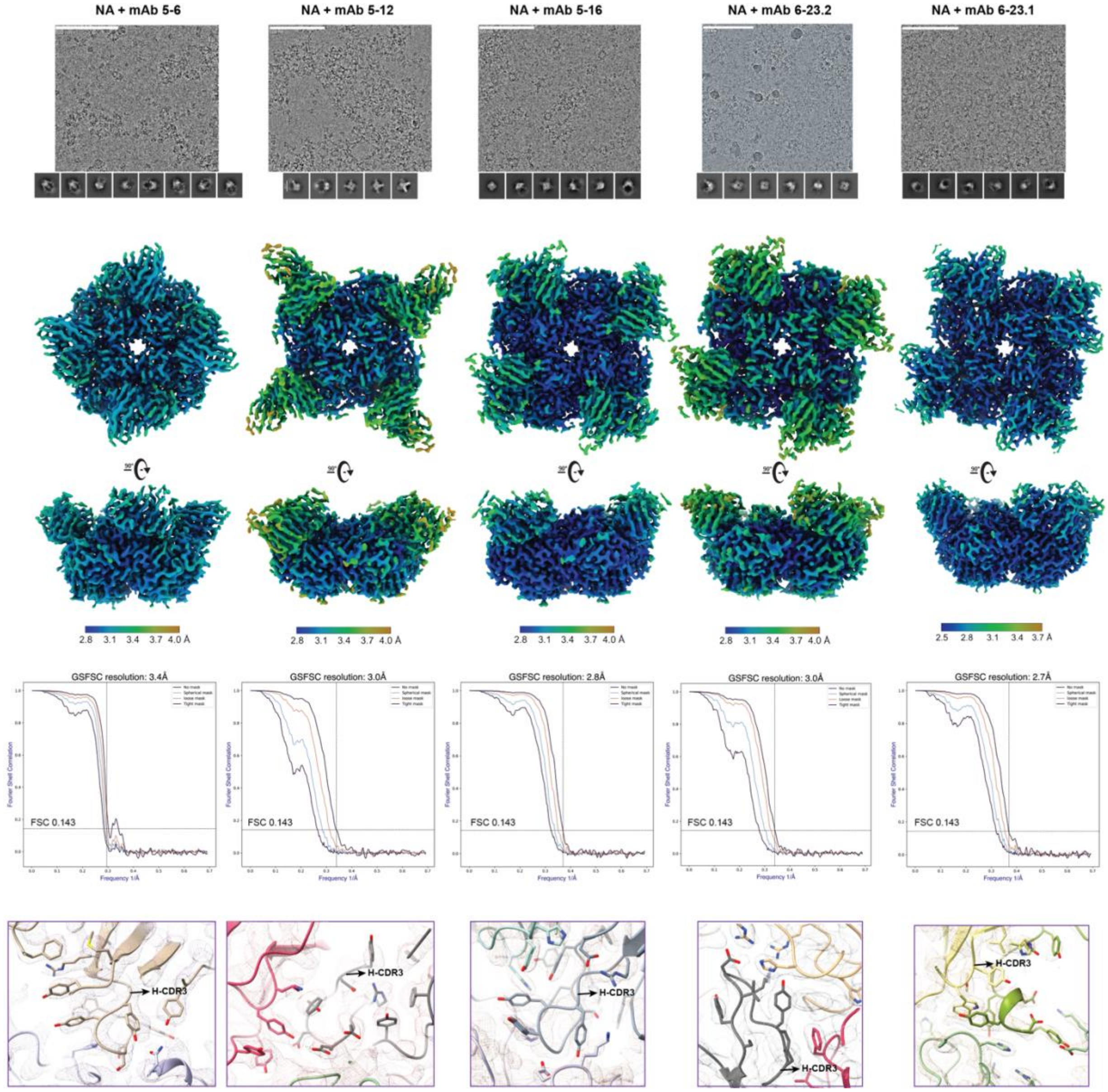


Supplemental Figure S7. Representative micrographs, 2D class averages, local resolution maps, FSC curves, and CDRH3s with electron potential maps for each NA-inhibiting mAb.

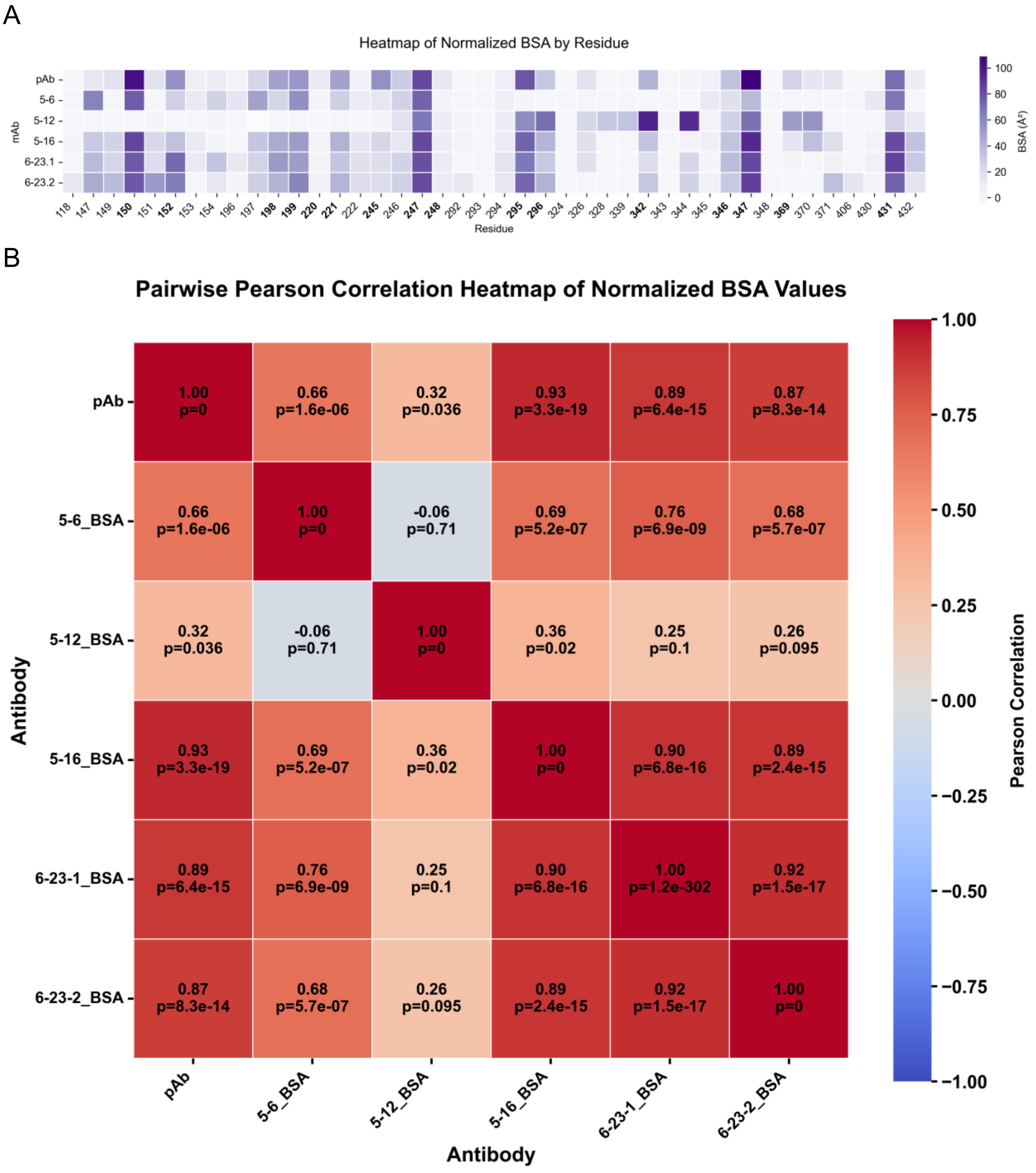


Supplemental Figure S8. A. Normalized heat map of antibody contributions to BSA of pAb vs the five isolated mAbs. Residues in bold correspond to pAb interaction residues. B. Pearson correlation coefficient of each mAb BSA per residue.
